# Zoledronate reduces loading-induced microdamage in cortical ulna of ovariectomized rats

**DOI:** 10.1101/2023.05.12.540579

**Authors:** Bohao Ning, Irène Londono, Catherine Laporte, Isabelle Villemure

## Abstract

As a daily physiological mechanism in bone, microdamage accumulation dissipates energy and helps to prevent fractures. However, excessive damage accumulation might bring adverse effects to bone mechanical properties, which is especially problematic among the osteoporotic and osteopenic patients treated by bisphosphonates. Some pre-clinical studies in the literature applied forelimb loading models to produce well-controlled microdamage in cortical bone. Ovariectomized animals were also extensively studied to assimilate human conditions of estrogen-related bone loss. In the present study, we combined both experimental models to investigate microdamage accumulation in the context of osteopenia and zoledronate treatment. Three-month-old normal and ovariectomized rats treated by saline or zoledronate underwent controlled compressive loading on their right forelimb to create *in vivo* microdamage, which was then quantified by barium sulfate contrast-enhanced micro-CT imaging. Weekly *in vivo* micro-CT scans were taken to evaluate bone (re)modeling and to capture microstructural changes over time. After sacrifice, three-point-bending tests were performed to assess bone mechanical properties. Results show that the zoledronate treatment can reduce cortical microdamage accumulation in ovariectomized rats, which might be explained by the enhancement of several bone structural properties such as ultimate force, yield force, cortical bone area and volume. The rats showed increased bone formation volume and surface after the generation of microdamage, especially for the normal and the ovariectomized groups. Woven bone formation was also observed in loaded ulnae, which was most significant in ovariectomized rats. Although all the rats showed strong correlations between periosteal bone formation and microdamage accumulation, the correlation levels were lower for the zoledronate-treated groups, potentially because of their lower levels of microdamage. The present study provides insights to further investigations of pharmaceutical treatments for osteoporosis and osteopenia. The same experimental concept can be applied in future studies on microdamage and drug testing.

## 1 Introduction

Bone microscopic damage (microdamage) accumulation is a physiological process that takes place during our daily activities (Burr et al., 1985; Frost, 1960). A balanced state of microdamage accumulation and repair can help to dissipate energy and to prevent fragility fractures (Diab et al., 2006; Diab & Vashishth, 2005). However, excessive levels of microdamage could lead to impaired bone mechanical properties (Burr et al., 1998; Pattin et al., 1996; Zioupos, 2001). Past studies in the literature have investigated how microdamage is initiated (Carter et al., 1981; Fazzalari et al., 1998; Nagaraja et al., 2005; Wenzel et al., 1996), accumulated (Diab & Vashishth, 2007; Thurner et al., 2006; Vashishth et al., 2000), propagated (O’Brien et al., 2005; Vashishth et al., 2003; Wang & Niebur, 2006), and repaired (Bentolila et al., 1998; Burr, 2002; Mori & Burr, 1993; Verborgt et al., 2000). Pre-clinical studies applying mechanical loading on long bones of small laboratory animals showed the advantage of creating consistent damage levels that can be quantified by histology (Bentolila et al., 1998; Herman et al., 2010; Seref-Ferlengez et al., 2014) or micro-CT-based methods (Gargac et al., 2014; Turnbull et al., 2011; Zhang et al., 2018).

Some experimental studies on microdamage made links with osteoporosis and osteopenia (Coates & Silva, 2020; Fazzalari et al., 1998; Lambers et al., 2013) because they can influence damage accumulation. These pathologies are characterized by low bone mineral density (BMD) (Dimai, 2017; NIH, 2001), which is especially common among the female post-menopausal population due to estrogen deficiency (Li & Xu, 2020; WHO, 1994). Ovariectomized rodent models were widely used in pre-clinical studies due to their strong resemblance to the human conditions of estrogen-related bone loss (Iwaniec & Turner, 2013; Jee & Yao, 2001; Kalu, 1991; Lelovas et al., 2008). As one of the common treatments for bone loss, various types of bisphosphonates (BPs) were tested regarding their effects on bone quality improvement (Burr, 2016; Feher et al., 2010; Hao et al., 2015; Kettenberger et al., 2014; Liu et al., 2017). Zoledronate (zoledronic acid) is considered among the most potent BPs by inhibiting the bone resorption thus increasing the BMD (Åstrand et al., 2006; Gasser et al., 2008; Kettenberger et al., 2014).

However, the BP-reduced bone resorption can decrease the efficiency of microdamage repair (Allen & Burr, 2007; Chapurlat & Delmas, 2009; Kidd et al., 2011; Li et al., 2001), leading to increased damage density (Allen, 2018; Ma et al., 2017; Mashiba et al., 2000) and therefore impaired bone quality (Allen & Burr, 2011; Burr, 2016, 2020). Clinical studies revealed that long term treatment with BPs can be associated with increased risk of atypical femoral fractures (Black et al., 2018; Black et al., 2021; Gedmintas et al., 2013; Shane et al., 2014), atypical forearm fractures (Moon et al., 2013; Oh et al., 2018; Ohta et al., 2022; Tan et al., 2015), and osteonecrosis of the jaw (Khosla et al., 2007; Marx, 2003; Shane et al., 2013). Although numerous translational studies were carried out to better understand these side effects (Kettenberger et al., 2014; Kidd et al., 2011; Mulcahy et al., 2015; Wang et al., 2019), the data are still lacking in terms of the direct relationships between microdamage accumulation, remodeling, and mechanical properties in the context of BP treatment. To fill in this knowledge gap, animal models are especially appropriate as they provide well-controlled experimental conditions.

In the present study, we combined the widely used ulna microdamage model with the well-established ovariectomized rat model to study microdamage accumulation in cortical ulna. Our objectives were: (1) to create and quantify *in vivo* ulnar microdamage in ovariectomized rats further treated by saline or zoledronate; (2) to quantify ulnar (re)modeling and mechanical properties; (3) to evaluate the correlations between ulnar microdamage and (re)modeling or mechanical parameters. We hypothesized that the zoledronate treatment can reduce loading-induced microdamage by improving structural properties in cortical ulna of ovariectomized rats.

## 2 Material and Methods

### 2.1 Experimental design

Three-month-old female virgin CD IGS rats (n=48) were received from the Charles River Laboratories on week 1 (Fig. 1). The animals were housed in cages of two with free access to food and water at the animal facility of the Sainte-Justine University Hospital Center. They lived a light/dark circle of 12 hours at about 23℃ and 21% humidity. One week of acclimation was allocated before the start of the experiment on week 2. The rats were assigned to one of the following treatment groups: “NOM” (normal, n=16), “OVX” (ovariectomized, n=16) and “ZOL” (zoledronate-treated, n=16). The NOM rats did not receive any surgery, serving as baseline controls, while the OVX and ZOL rats received bilateral ovariectomy at the animal supplier on week 0 (Fig. 1). Within each treatment group (n=16/group), the rats were randomly assigned to two sub-groups of different sacrifice time points (n=8 at week 11 and n=8 at week 12). The effects of ovariectomy have been confirmed, with significant bone loss occurring in trabecular tibia from week 2 to week 12 (Supplementary Material). The Institutional Committee for Good Animal Practices in Research of the Sainte-Justine University Hospital Center approved the animal experiments in this study.

**Fig. 1.**
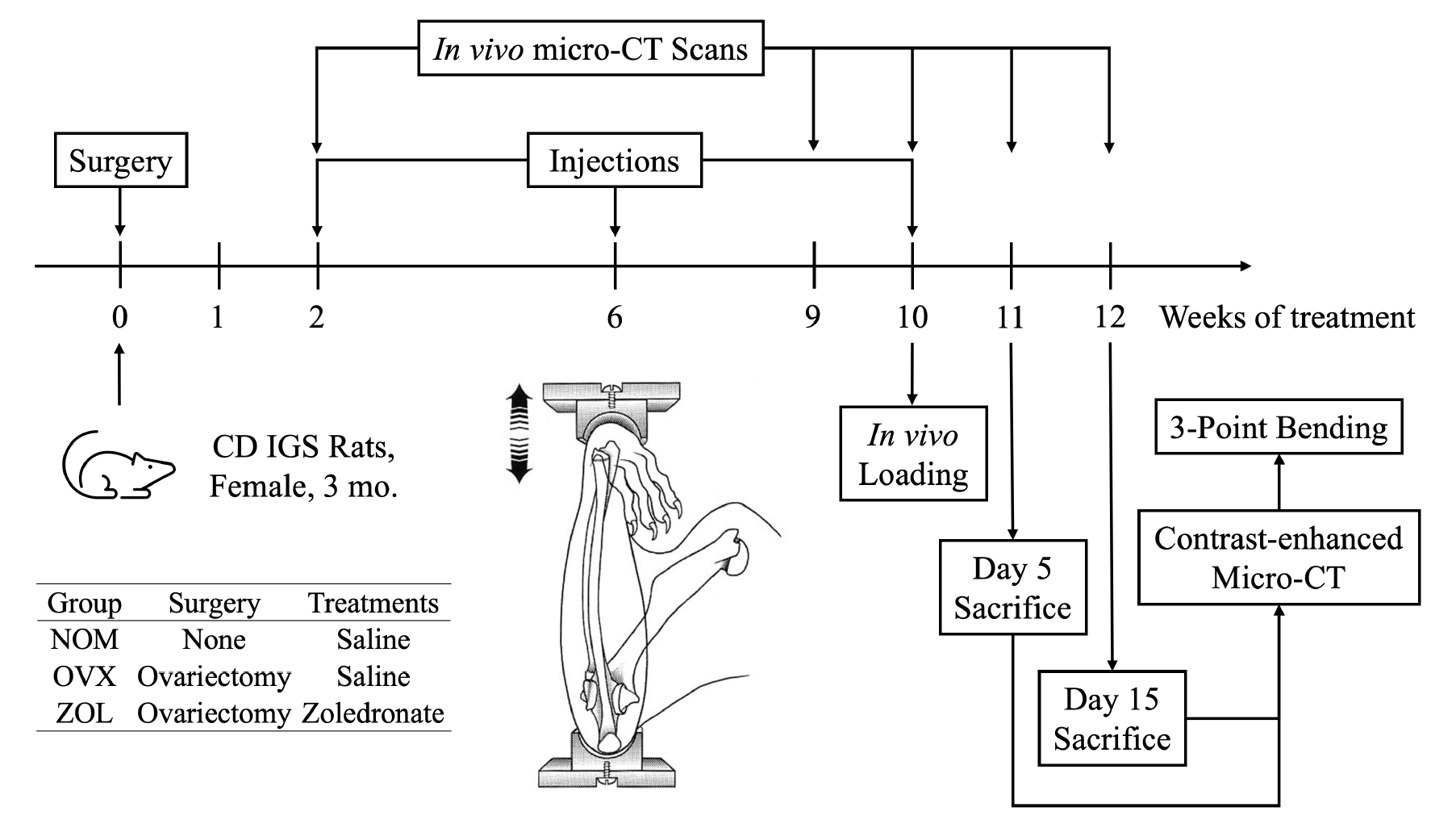
Experimental design. The forelimb loading diagram was adapted from (Robling et al., 2002) with permission obtained from the publisher.

As shown in Fig. 1, the rats received *in vivo* micro-CT scans on weeks 2, 9, 10, 11 and 12 to follow up on the microstructural changes in their forelimbs and hindlimbs (section 2.2). They also received subcutaneous injections on weeks 2, 6 and 10, where the NOM and OVX rats were given sterile saline (1 mL/kg) and the ZOL rats were given zoledronate (100 μg/kg) for a total treatment period of 9 weeks. The dose of the zoledronate was based on past rodent models to reflect the clinical doses used for bone loss treatment in humans (Aref et al., 2016; Feher et al., 2010; Gasser et al., 2008). On week 10, all the rats received a single session of mechanical loading (section 2.3) on their right forelimb to generate microdamage *in vivo*. Half of the rats (n=8 from each treatment group) were sacrificed 5 days post-loading (week 11), and the other half 15 days post-loading (week 12) to quantify the amount of microdamage using contrast-enhanced micro-CT imaging (section 2.4). After sacrifice, all the ulnar samples received a three-point-bending test to evaluate their mechanical properties (section 2.5).

### 2.2 *In vivo* micro-CT imaging

On week 2, *in vivo* micro-CT scans were acquired using a SkyScan 1176 microtomograph (Bruker microCT N.V., Kontich, Belgium) on rats’ right forelimb (mid-diaphysis) to capture the initial bone microstructure (Fig. 1). On weeks 9, 10, 11 and 12, additional weekly scans were performed on both sides of forelimb to calculate the (re)modeling rates. Rod-shaped phantoms of 0.25 and 0.75 g/cm^3^ were scanned as well for calibration purposes. All the scans were performed at an image resolution of 8.74 μm, using a filter of 1 mm Al, exposure time of 1200 ms, voltage of 65 kVp, current of 300 μA, rotation angle of 0.5°/180°. The rats were under isoflurane anesthesia (2.5%) supplied by oxygen (1.5 L/min) during the scans, which lasted about 11 minutes each. The radiation exposure was estimated to be about 1.08 Gy for each scan by the CTion software (Bruker microCT N.V., Kontich, Belgium) (Mustafy et al., 2018).

After the scans, the images were reconstructed using the NRecon software (Bruker microCT N.V., Kontich, Belgium). The dynamic range was set to [0, 0.06] (attenuation coefficient) and the post-alignment was adjusted visually starting from the default value proposed by the software. Beam-hardening correction was applied at 20% and ring-artifact correction was set to 5. For the subsequent image analysis, volumes of interest (VOIs) were extracted using CT Analyzer (Bruker microCT N.V., Kontich, Belgium). The ulnar VOIs were centered at the mid-third diaphysis, expanding for about 20% of total bone length. This location was shown to receive the most flexion in response to axial forelimb compression (Herman et al., 2010; Zhang et al., 2018). A global threshold of 75/255 (0.45 g/cm^3^) was set for the following image analysis, which was determined by an automatic thresholding function in CT Analyzer and visually verified in the images and histograms.

To calculate the weekly (re)modeling parameters between weeks 9 and 12, the ulnar scans from weeks 10, 11 and 12 were rigidly registered to the baseline scans of week 9 using custom software developed using ITK (Kitware Inc., Clifton Park, NY, USA) (Johnson et al., 2015). Automatic registration was based on a regular step gradient optimizer with mutual information as the optimization metric. Once the images were registered, a previously validated MATLAB-based program was used to calculate the (re)modeling parameters on the periosteum (Birkhold et al., 2014). The following (re)modeling parameters were analyzed in terms of volume, surface, and thickness: newly mineralized bone volume per bone volume (MV/BV), newly eroded bone volume per bone volume (EV/BV), newly mineralized bone surface per bone surface (MS/BS), newly eroded surface per bone surface (ES/BS), mineral apposition rate (MAR, μm/day), and mineral resorption rate (MRR, μm/day). The static histomorphometry was performed using CT Analyzer (Birkhold et al., 2016; Bouxsein et al., 2010). The following cortical ulnar parameters were evaluated: tissue mineral density (TMD, g/cm^3^), cortical bone volume (Ct.BV, mm^3^), average cross-sectional cortical area (Ct.Ar, mm^2^), and average cortical thickness (Ct.Th, μm).

### 2.3 *In vivo* mechanical loading

The *in vivo* loading session was applied on week 10 (Fig. 1). One hour before loading, the rats were given a dose of buprenorphine (0.05 mg/kg) subcutaneously as a sedative and an analgesic. Ten minutes before loading, the rats were put into anesthesia via isoflurane (3.0%) supplied by oxygen (1.8 L/min). They were laid on their back, supported by a surface warming pad (36 to 37°C). The *in vivo* loading, performed using a Mach-1 micromechanical system (Biomomentum Inc., Laval, QC, Canada), started with a preconditioning session of 50 sinusoidal cycles of cyclic compression with a frequency of 2 Hz and a displacement amplitude of 1 mm.

The subsequent main loading step was based on sinusoidal cyclic compression as well with a physiological frequency of 2 Hz and a peak force of 16 N, leading to a cumulative actuator displacement of 0.4591 mm and 4665 loading cycles on average. The loading was stopped when one of the following equivalent conditions was satisfied: the cumulative actuator displacement reached 0.60 mm, or 5040 cycles was completed. The dual stopping criteria served to prevent the forelimb from fracture, which corresponded to 58% of the fracture point (1.0391 mm of the cumulative actuator displacement) based on a preliminary test where 90% of the rats (n=20 in total) did not encounter a fracture within 0.60 mm of the cumulative actuator displacement (equivalent to 5040 cycles). Right after the *in vivo* loading, the rats were given a dose of metacam (1 mg/kg) subcutaneously to relieve potential pain caused by loading. The rats were followed up 1h, 4h, 24h, 48h and 72h after the loading session for assessment of their well-being, based on which no additional analgesic was provided.

### 2.4 BaSO_4_ Staining and contrast-enhanced micro-CT imaging

Once the rats were sacrificed by CO_2_ asphyxiation and decapitation, the forelimbs were collected, removed of soft tissues, and dissected into individual ulnar samples. The samples were wrapped in phosphate-buffered-saline (PBS)-soaked gauze and stored in tubes of 5 mL in a freezer (−20℃). Five days after the dissection, the samples were thawed for barium sulfate (BaSO_4_) contrast staining according to a previously validated method (Choudhari et al., 2016; Gargac et al., 2014; Landrigan et al., 2011; Leng et al., 2008; Turnbull et al., 2011; Zhang et al., 2018): the ulnar samples were immersed in an equally mixed solution of 0.5 M barium chloride (BaCl_2_), acetone, and PBS solution for 3 days under vacuum; followed by another 3 days of staining in an equally mixed solution of 0.5 M sodium sulfate (Na_2_SO_4_), acetone, and PBS solution under vacuum.

After the staining, the samples were gently washed and wrapped in PBS-soaked gauze for contrast-enhanced micro-CT imaging. The scans were performed using the SkyScan 1176 at an image resolution of 8.74 μm, using a filter of 1 mm Al, exposure time of 1500 ms, voltage of 65 kVp, current of 385 μA, rotation angle of 0.25°/180°. After the scans, the samples were wrapped again in PBS-soaked gauze and put back into tubes for storage (−20℃). The micro-CT images were reconstructed using the NRecon software. The dynamic range was set to [0, 0.12] (attenuation coefficient) and the post-alignment was adjusted visually starting from a default value proposed by the software. Beam-hardening correction was applied at 20% and ring-artifact correction was set to 5.

To evaluate the microdamage accumulated in ulna, the same VOIs were used as explained in section 2.2. The microdamage was intended to be evaluated inside the original pre-loading bone volume. Therefore, the newly formed woven bone on the periosteum was excluded by registering the ulnar VOIs to the images of week 10 (pre-loading images). The image registration method was the same as explained in section 2.2. Subsequently, the VOIs were imported into MATLAB R2018b (The MathWorks, Inc., Natick, MA, USA) for microdamage quantification. A customized script evaluated the stained volume over the total bone volume (SV/BV), which reflects the overall extent of microdamage based on methods developed in previous studies, which were validated by the histology-based methods (Turnbull et al., 2011; Zhang et al., 2018). The thresholds for the bone volume and the stained volume were 0.67 g/cm^3^ and 1.77 g/cm^3^ respectively, which were determined based on histogram analysis for each of the 3D datasets. The threshold for the stained volume has also been verified to be well above the dynamic range of the bone mineral density. For each pair of contralateral ulnae, the difference in SV/BV between the loaded and the control samples was calculated to derive a new parameter ΔSV/BV, which reflects the overall amount of microdamage resulting from the applied *in vivo* loading, excluding the effects of daily damage as well as staining in the voids and on the bone surface. Only one pair of samples (2%) showed slightly higher SV/BV on the control sample than on the loaded sample. In this case, we took the value of ΔSV/BV to be 0 and interpreted the result as insignificant damage produced by the *in vivo* loading.

### 2.5 Three-point-bending

One week after the contrast-enhanced micro-CT, the ulnar samples were thawed to room temperature (about 23℃) for the three-point-bending procedure. The bending was performed using the Mach-1 micromechanical system and a customized bending platform. The span between the two sample supports was set to 18 mm. The samples were loaded in the medio-lateral direction (Fig. 2) to ensure stable positioning of the samples. The actuator displacement speed was set to 0.01 mm/s and the data acquisition was performed at a frequency of 100 Hz.

**Fig. 2.**
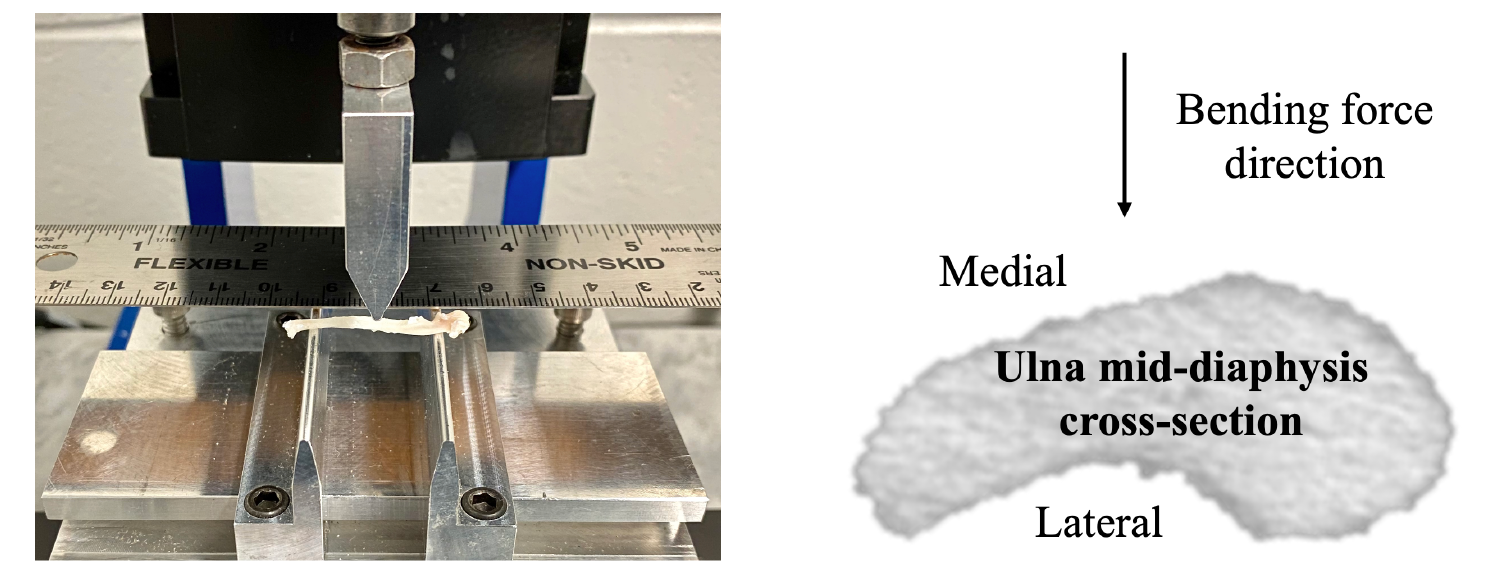
Three-point-bending setup for the ulna samples. The experimental setup is shown on the left and the cross-sectional view is shown on the right.

After the bending tests, the force-displacement curves were analyzed using the Mach-1 Analysis software (Biomomentum Inc., Laval, QC, Canada) to measure the following structural properties: ultimate force (F_u_, N), yield force (F_y_, N), stiffness (S, N/mm), pre-yield displacement (D_pre_, mm), and pre-yield energy (U_pre_, mJ). To estimate the material (tissue) properties using beam theory equations (Burr, 2020; Burr et al., 2015; Mashiba et al., 2000; Schriefer et al., 2005; Wallace, 2019), the *ex vivo* micro-CT images of the samples were used to calculate the second moment of area, maximum diameter of the loaded section, and the distance from the neutral axis to the bone surface with the BoneJ plugin (Domander et al., 2021) of the FIJI software (Schindelin et al., 2012). The following material properties were calculated: ultimate stress (σ_u_, MPa), yield stress (σ_y_, MPa), Young’s modulus (E, GPa), pre-yield strain (ɛ_pre_, μɛ) and pre-yield toughness (u_pre_, MPa). The post-yield properties were not analyzed as the conversion equations are only applicable in the elastic region (Wallace, 2019).

### 2.6 Statistics

The statistical tests were performed using SPSS Statistics v26 (IBM Corp, Armonk, NY, USA). Different ANOVA tests were performed for each parameter as shown in Table 1. The assumptions of the ANOVA tests were verified when applicable. The boxplots revealed very few outliers (less than 3%) which were not removed since they did not consist of data entry or measurement errors to our knowledge. The Shapiro-Wilk’s test and Normal Q-Q Plots determined that the data were normally distributed in general. The Levene’s test for equality of variances was performed, showing that variances were homogenous for the vast majority of the data groups. The ANOVA test was performed because it is somewhat robust to heterogeneity when the group sample sizes are equal, data are normal, and the ratio of the largest group variance to the smallest group variance is less than 3 (Altman, 1991; Jaccard, 1998; Keppel, 2004; Laerd Statistics, 2015). The Box’s test determined that the homogeneity of covariances was also met for most of the data groups. As determined by the Mauchly’s test of sphericity, the assumption of sphericity was violated for some data groups. Thus, the Greenhouse–Geisser correction was applied when interpreting the within-subject effects. Bonferroni correction was applied for the *p*-value calculation in the *post hoc* tests and multiple comparisons, after which the statistical significance level (alpha) was set to 0.05. Additional data of interest are reported for the results of bone mechanical properties (*p* < 0.10).

**Table 1.** Summary of ANOVA tests performed. The treatment factor refers to the rat group NOM, OVX and ZOL. The recovery factor refers to the sacrifice time point of day 5 or day 15 post-loading. The bone side factor refers to left (control) or right (loaded) ulna. The time points vary from week 2 to week 12 of the experiment.

| Parameter | ANOVA Type | Between-subjects Factors | Within-subjects Factor |
| --- | --- | --- | --- |
| Bone microdamage | Two-way | Treatment, recovery | n/a |
| Bone (re)modeling | Three-way mixed | Recovery, bone side | Time Points |
| Bone mechanical properties | Three-way | Treatment, recovery, bone side | n/a |
| Static histomorphometry | Two-way mixed | Treatment | Time Points |

Besides the ANOVA analyses, two-tailed t-tests were performed for the static histomorphometry parameters on week 12 to compare between left (control) and right (loaded) ulna of the same rat. The statistical significance level (alpha) was set to 0.05 for all the t-tests. The two-tailed Spearman’s rank-order correlation was used to evaluate the correlations of the bone microdamage parameter SV/BV with bone (re)modeling or mechanical properties. The analysis between SV/BV and the (re)modeling parameters was only performed for the rats that were sacrificed 15 days post-loading, since the (re)modeling parameters of the day-5 rats were not available. The assumption of a monotonic relationship between all the data pairs was verified using scatterplots. The Spearman’s correlation coefficient was considered significant for *p* < 0.05, and additional data of interest (*p* < 0.15) are reported as well.

## 3 Results

### 3.1 Bone microdamage

An example visualization of ulnar microdamage is shown in Fig. 3, representative of each treatment group and the side of bone. The volume of formed, resorbed and constant bone were identified based on the registration results between the stained VOIs (week 12) and the pre-loading VOIs (week 10). The damage (stained) volume was identified based on the threshold analysis of the stained VOIs. The results of ΔSV/BV are shown in Fig. 4. Both NOM and OVX rats had increased ΔSV/BV over time; the ZOL ulnae were at a lower level on day 15 compared to the OVX. The inter-group differences of ΔSV/BV were most significant at week 12. Therefore, the example visualization in Fig. 3 focuses on the data from week 12.

**Fig. 3.**
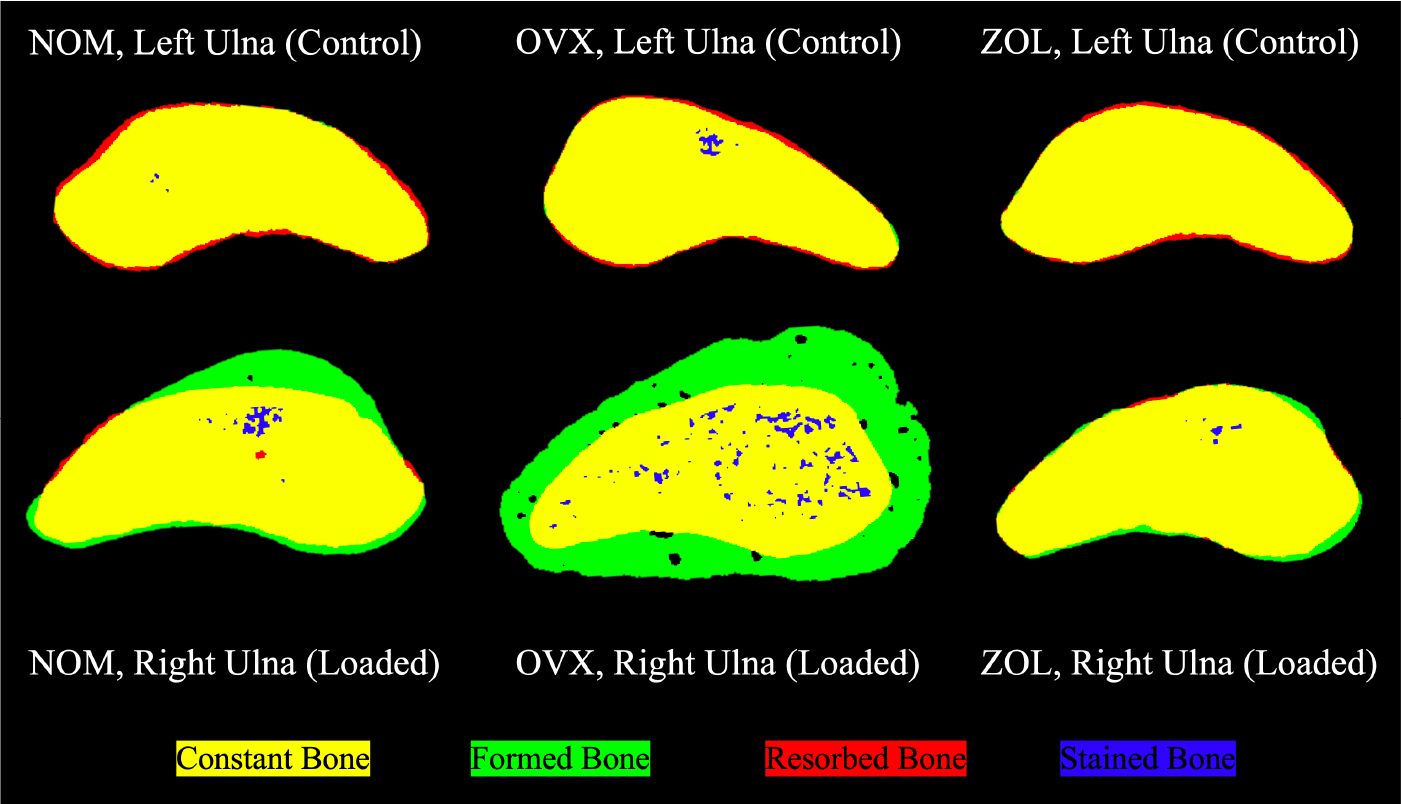
Example visualization of the ulna microdamage results shown in binarized cross-sectional area. The top row shows the control ulnae of each group, and the bottom row shows the loaded ulnae. Based on the image registration results and threshold analysis, the constant, formed, resorbed, and stained bone volume are shown in yellow, green, red, and blue, respectively.

**Fig. 4.**
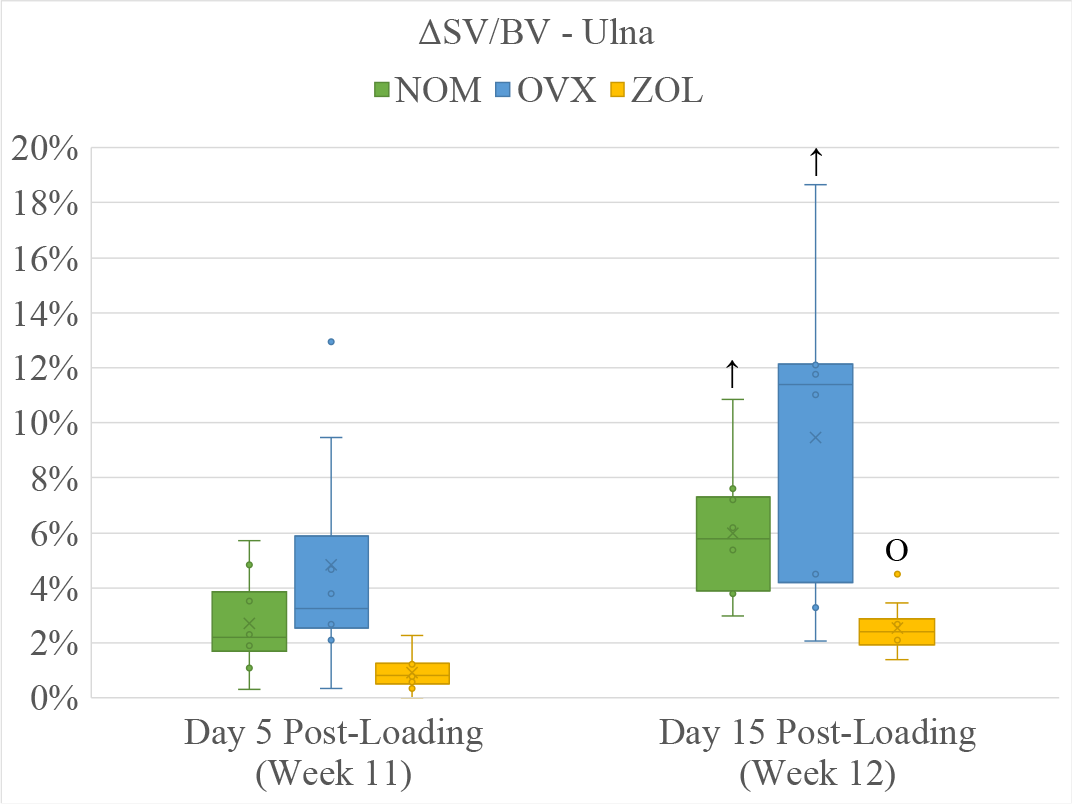
ΔSV/BV for ulna, reflecting the overall microdamage accumulated by *in vivo* loading (n=8/group). “↑” refers to a significant increase compared to the previous time point. “O” refers to a significant difference compared to the OVX group. Bonferroni correction was used for *post hoc* tests and multiple comparison adjustments, after which the statistical significance level (alpha) was set to 0.05.

### 3.2 Bone (re)modeling

The results of (re)modeling are shown in Fig. 5. Effects of recovery time were observed for almost all the parameters. Effects of treatment groups were apparent as well. For example, the MV/BV and MS/BS showed approximatively three levels on week 12: the NOM (loaded) and OVX (loaded) on a higher level, the NOM (control) and ZOL (loaded) on the middle level, then the OVX (control) and ZOL (control) on a relatively lower level. In general, the resorption parameters showed less between-group differences than their formation counterparts. Contralateral differences of each parameter were not always clear, and statistically significant results were mostly in week 12.

**Fig. 5.**
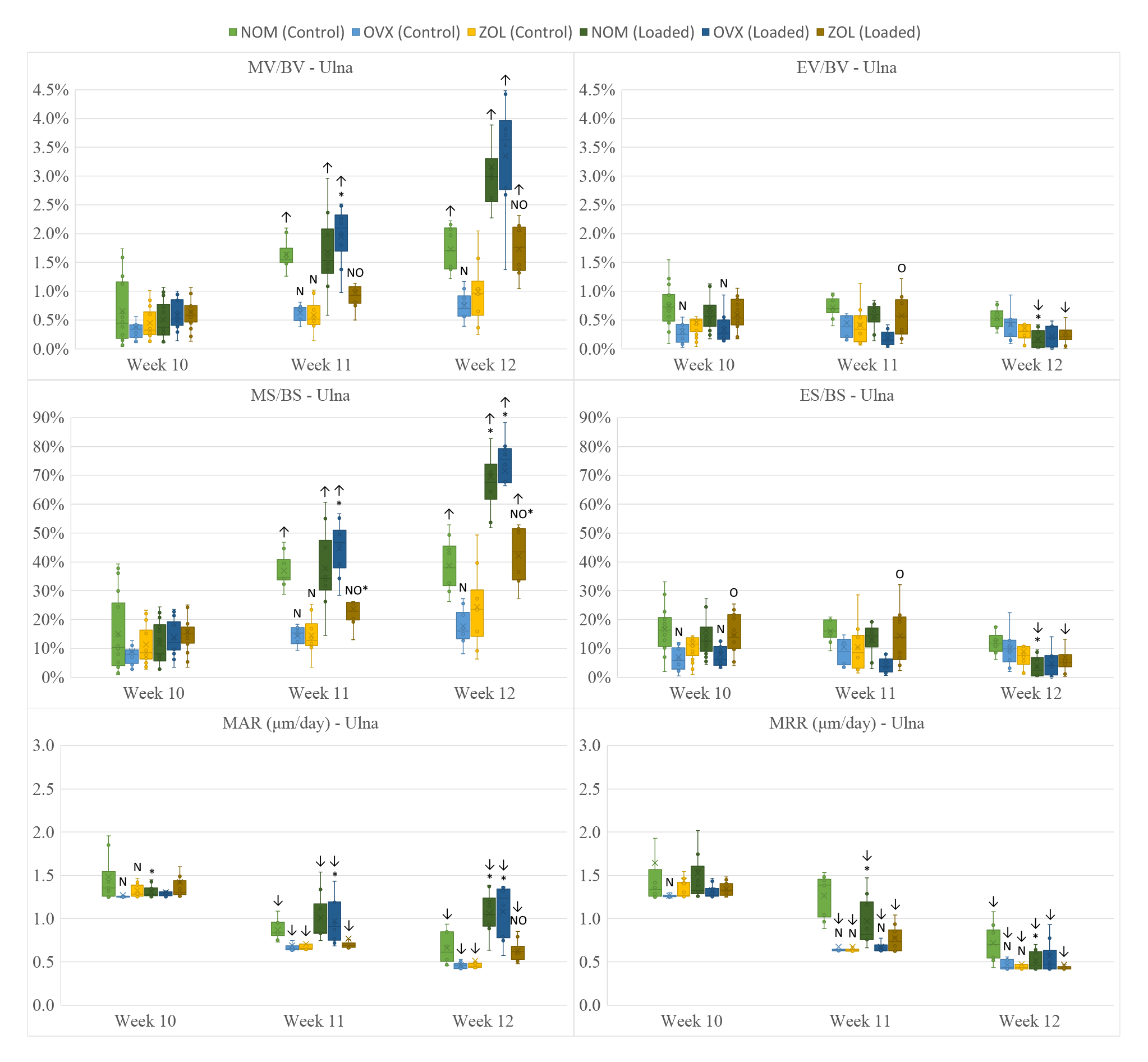
Ulnar (re)modeling parameters shown in boxplots (n=16/group at week 10, n=8/group at weeks 11 and 12). “↑” or “↓” refers to a significant increase or decrease compared to week 10. “N” or “O” refers to a significant difference compared to the NOM or OVX group. “*” refers to a significant difference between the left and right ulna. Bonferroni correction was applied for the *p*-value calculation in the *post hoc* tests and multiple comparisons, after which the statistical significance level (alpha) was set to 0.05.

### 3.3 Bone mechanical properties

The results of the mechanical properties (structure and material) are shown in Table 2. Effects of recovery time were observed for the NOM and OVX ulnae. The ZOL ulnae showed higher F_u_, F_y_ and S compared to the NOM or OVX ulnae at most of the relevant data points. Their material counterparts σ_u_, σ_y_ and E showed less consistent results. There were a few data points showing contralateral differences (control vs loaded), which were mostly found in results of loaded samples on day 15 post-loading.

**Table 2.** Ulnar mechanical properties tested by three-point-bending (n=8/group). The structural properties are shown on the top half of the chart, and their material counterparts are at the bottom half. “↑” or “↓” refers to a significant increase or decrease compared to day 5. “N” or “O” refers to a significant difference compared to the NOM or OVX group. “*” refers to a significant difference between the left (control) and right (loaded) ulna. Bonferroni correction was applied for the *p*-value calculation in the *post hoc* tests and multiple comparisons, after which the statistical significance level (alpha) was set to 0.05. The result of a *p*-value between 0.05 and 0.10 was marked in blue. The cells of statistical significance are in bold.

| Time | VOI | Group | $F_u$ (N) | $F_y$ (N) | $S$ (N/mm) | $D_{pre}$ (mm) | $U_{pre}$ (mJ) |
| --- | --- | --- | --- | --- | --- | --- | --- |
| Day 5 | Control | NOM | 11.68 ± 1.37 | 9.06 ± 0.87 | 17.54 ± 2.09 | 0.5957 ± 0.0636 | 2.67 ± 0.33 |
|  |  | OVX | 13.41 ± 1.86 | 10.34 ± 1.94 | **26.23 ± 5.66** N | 0.5366 ± 0.0686 | 2.70 ± 0.72 |
|  |  | ZOL | **16.42 ± 3.58** NO | **11.47 ± 2.12** N | **30.05 ± 13.55** N | 0.5542 ± 0.1359 | 3.10 ± 1.02 |
|  | Loaded | NOM | 11.94 ± 1.47 | 9.43 ± 1.23 | 17.70 ± 1.71 | 0.6159 ± 0.0559 | 2.87 ± 0.57 |
|  |  | OVX | 12.69 ± 2.32 | 10.26 ± 1.98 | 25.09 ± 8.33 | 0.5774 ± 0.1261 | 2.83 ± 0.68 |
|  |  | ZOL | **16.63 ± 3.50** NO | **11.84 ± 2.66** N | **26.59 ± 7.16** N | 0.6056 ± 0.1025 | 3.43 ± 1.16 |
| Day 15 | Control | NOM | 12.93 ± 2.34 | 10.08 ± 1.62 | 22.33 ± 5.84 | 0.6437 ± 0.0886 | 2.87 ± 0.48 |
|  |  | OVX | 12.47 ± 1.38 | 9.37 ± 0.87 | **20.20 ± 3.94** ↓ | 0.5852 ± 0.0894 | 2.63 ± 0.30 |
|  |  | ZOL | **16.65 ± 2.22** NO | **11.72 ± 1.68** O | **31.39 ± 5.58** NO | **0.5467 ± 0.0813** N | 2.99 ± 0.66 |
|  | Loaded | NOM | **14.22 ± 1.61** ↑ | 10.39 ± 1.19 | **24.38 ± 3.69** ↑ | **0.5578 ± 0.0816** * | 2.88 ± 0.62 |
|  |  | OVX | 14.23 ± 2.22 | 9.87 ± 1.58 | **26.47 ± 7.04** * | **0.4741 ± 0.0448** ↓* | 2.33 ± 0.44 |
|  |  | ZOL | 16.47 ± 2.11 | **11.84 ± 1.72** O | 28.96 ± 4.80 | 0.5381 ± 0.0310 | **3.14 ± 0.57** O |
| Time | VOI | Group | $\sigma_u$ (MPa) | $\sigma_y$ (MPa) | $E$ (GPa) | $\epsilon_{pre}$ (µε) | $u_{pre}$ (MPa) |
| Day 5 | Control | NOM | 275.63 ± 14.58 | 214.29 ± 12.83 | 22.53 ± 2.10 | 10,998 ± 1,477 | 1.17 ± 0.15 |
|  |  | OVX | 262.79 ± 20.45 | 202.53 ± 28.18 | 25.42 ± 3.07 | 10,718 ± 1,034 | 1.06 ± 0.26 |
|  |  | ZOL | 290.13 ± 13.96 | 205.55 ± 26.36 | 24.67 ± 5.84 | 11,326 ± 2,368 | 1.16 ± 0.36 |
|  | Loaded | NOM | 262.81 ± 22.70 | 207.34 ± 15.50 | 20.23 ± 1.83 | 11,910 ± 1,291 | 1.21 ± 0.18 |
|  |  | OVX | 243.61 ± 28.97 | 196.84 ± 25.94 | 23.80 ± 6.39 | 11,620 ± 2,732 | 1.10 ± 0.27 |
|  |  | ZOL | **288.54 ± 43.35** O | 205.64 ± 34.34 | 21.88 ± 3.86 | 12,648 ± 2,102 | 1.25 ± 0.37 |
| Day 15 | Control | NOM | 289.05 ± 22.83 | 214.10 ± 15.01 | 22.41 ± 2.32 | **13,507 ± 928** ↑ | **1.42 ± 0.18** ↑ |
|  |  | OVX | **250.86 ± 21.28** N | **188.50 ± 12.61** N | **21.12 ± 3.98** ↓ | **11,287 ± 1,805** N | **1.02 ± 0.14** N |
|  |  | ZOL | **283.42 ± 28.60** O | 199.72 ± 23.32 | 25.16 ± 3.72 | **11,608 ± 1,876** N | **1.08 ± 0.24** N |
|  | Loaded | NOM | 274.10 ± 18.20 | 203.02 ± 15.16 | 21.31 ± 1.51 | **13,618 ± 1,379** ↑ | 1.36 ± 0.20 |
|  |  | OVX | 253.65 ± 34.43 | **176.50 ± 29.40** N↓ | 22.63 ± 3.97 | **9,820 ± 1,043** N↓* | **0.87 ± 0.20** N↓ |
|  |  | ZOL | 268.66 ± 23.23 | 192.86 ± 16.92 | **21.97 ± 2.52** * | **11,551 ± 984** N | **1.10 ± 0.17** N |

### 3.4 Static histomorphometry

The results of static histomorphometry are shown in Table 3. The effects of treatment time were significant for the OVX and ZOL rats. The ZOL ulnae (control) had higher Ct.BV than the NOM, and higher Ct.Ar than both NOM and OVX rats on week 12. Only the NOM and OVX groups showed some significant differences between their control and loaded ulnae on week 12.

**Table 3.**
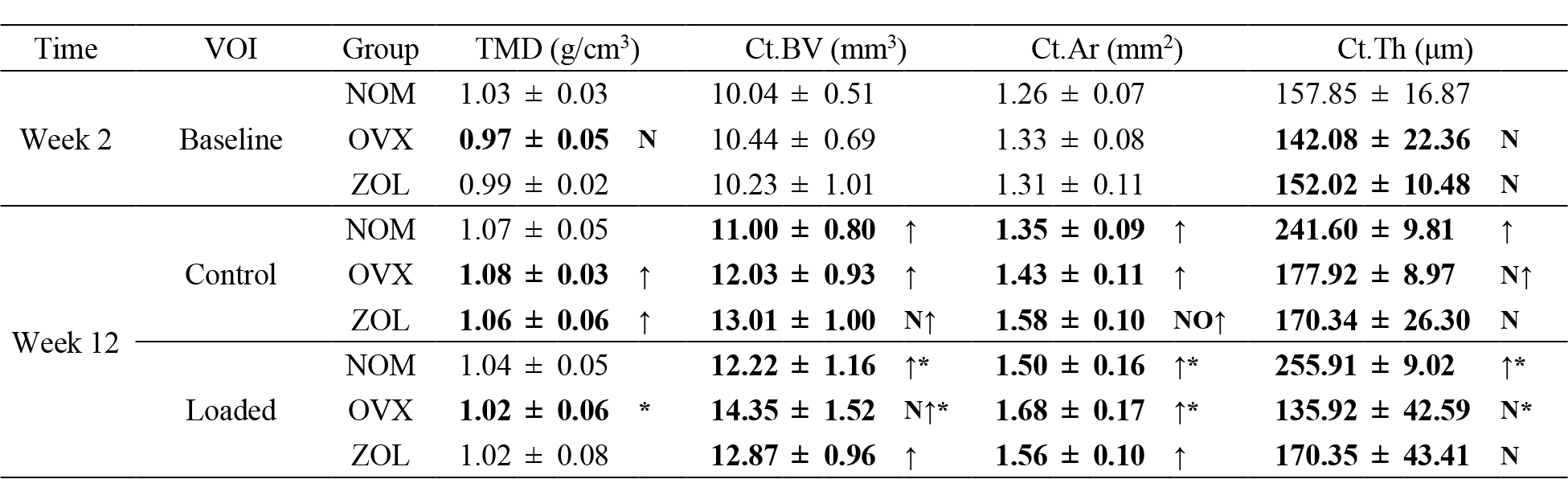
Ulnar static histomorphometry (n=8/group). “↑” refers to a significant increase compared to week 2. “N” or “O” refers to a significant difference compared to the NOM or OVX group. “*” refers to a significant difference between the left (control) and right (loaded) ulna. Bonferroni correction was applied for the *p*-value calculation in the *post hoc* tests and multiple comparisons, after which the statistical significance level (alpha) was set to 0.05. The cells of statistical significance are in bold.

| Time | VOI | Group | TMD (g/cm<sup>3</sup>) | Ct.BV (mm<sup>3</sup>) | Ct.Ar (mm<sup>2</sup>) | Ct.Th (μm) |
| --- | --- | --- | --- | --- | --- | --- |
| Week 2 | Baseline | NOM | 1.03 ± 0.03 | 10.04 ± 0.51 | 1.26 ± 0.07 | 157.85 ± 16.87 |
|  |  | OVX | **0.97 ± 0.05** N | 10.44 ± 0.69 | 1.33 ± 0.08 | **142.08 ± 22.36** N |
|  |  | ZOL | 0.99 ± 0.02 | 10.23 ± 1.01 | 1.31 ± 0.11 | **152.02 ± 10.48** N |
| Week 12 | Control | NOM | 1.07 ± 0.05 | **11.00 ± 0.80** ↑ | **1.35 ± 0.09** ↑ | **241.60 ± 9.81** ↑ |
|  |  | OVX | **1.08 ± 0.03** ↑ | **12.03 ± 0.93** ↑ | **1.43 ± 0.11** ↑ | **177.92 ± 8.97** N↑ |
|  |  | ZOL | **1.06 ± 0.06** ↑ | **13.01 ± 1.00** N↑ | **1.58 ± 0.10** NO↑ | **170.34 ± 26.30** N |
|  | Loaded | NOM | 1.04 ± 0.05 | **12.22 ± 1.16** ↑* | **1.50 ± 0.16** ↑* | **255.91 ± 9.02** ↑* |
|  |  | OVX | **1.02 ± 0.06** * | **14.35 ± 1.52** N↑* | **1.68 ± 0.17** ↑* | **135.92 ± 42.59** N* |
|  |  | ZOL | 1.02 ± 0.08 | **12.87 ± 0.96** ↑ | **1.56 ± 0.10** ↑ | **170.35 ± 43.41** N |

### 3.5 Correlation analysis

The correlation results between SV/BV and the (re)modeling parameters are shown in Table 4. All the formation parameters showed significant positive correlations with the microdamage parameter SV/BV. The correlations were mostly negative for the resorption parameters, and the NOM rats had statistically significant results for all the parameters. The correlations between SV/BV and the mechanical properties are shown in Table 5. All the data points with statistical significance showed negative correlations.

**Table 4.**
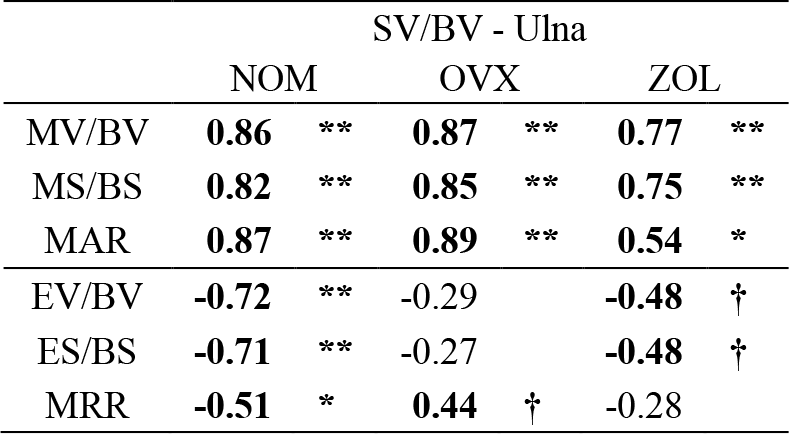
Spearman’s correlation coefficients between SV/BV and the (re)modeling parameters (n=8/group). The data are marked bold for statistical significance (“**” for *p* < 0.01, “*” for *p* < 0.05, and “†” for *p* < 0.15).

**Table 5.** Spearman’s correlation coefficients between SV/BV and the mechanical properties (n=16/group). The data are marked bold for statistical significance (“**” for *p* < 0.01, “*” for *p* < 0.05, and “†” for *p* < 0.15).

|  | SV/BV - Ulna |  |  |  |
| --- | --- | --- | --- | --- |
|  | NOM |  | OVX | ZOL |
| F<sub>u</sub> | 0.18 |  | -0.12 | **-0.26** † |
| F<sub>y</sub> | 0.03 |  | -0.21 | -0.21 |
| S | 0.16 |  | -0.13 | -0.19 |
| D<sub>pre</sub> | **-0.34** | † | -0.13 | -0.07 |
| U<sub>pre</sub> | -0.13 |  | -0.22 | -0.17 |
| σ<sub>u</sub> | **-0.30** | † | -0.23 | **-0.31** † |
| σ<sub>y</sub> | **-0.32** | † | -0.23 | **-0.27** † |
| E | **-0.39** | * | -0.19 | -0.24 |
| ε<sub>pre</sub> | 0.16 |  | -0.11 | -0.13 |
| u<sub>pre</sub> | 0.02 |  | -0.14 | -0.23 |

## 4 Discussion

### 4.1 The zoledronate treatment improves osteopenic conditions

Ovariectomy-induced changes on cortical bone (ulna, Table 3) were less obvious than on trabecular bone (tibia, Supplementary Material). Evidence from the literature suggests that bone loss in ovariectomized rats was quicker in trabecular bone than in cortical bone (Jee & Yao, 2001). It is also easier to detect porosity in trabecular bone than in cortical bone using laboratory micro-CT scanners (Cooper et al., 2007). Nonetheless, the ZOL rats had lower ΔSV/BV (Fig. 4) and higher F_u_, F_y_, S, σ_u_, Ct.Ar (at some data points in Table 2 and Table 3) compared to the OVX rats. The beneficial effects of zoledronate treatment were found significant for improving the bone strength (F_u_, σ_u_) without sacrificing the bone brittleness (D_pre_, ɛ_pre_) (Bilston et al., 2002). Past experimental studies showed that zoledronate treatment can improve the bending strength of long bones (Aref et al., 2016; Bilston et al., 2002), which was positively correlated with BMD (Hornby et al., 2003). Trabecular (Hornby et al., 2003) and cortical (Stadelmann et al., 2011) microstructure could also be improved by the zoledronate treatment. We confirm that zoledronate can improve the bone quality of ovariectomized rats in terms of bone microstructure and bending strength (Aref et al., 2016; Gasser et al., 2008; Meixner et al., 2017).

### 4.2 The zoledronate treatment reduces cortical microdamage accumulation

The ZOL rats accumulated less microdamage than the other rats as shown by the lower ΔSV/BV compared to the OVX ulnae on day 15 post-loading. In addition, levels of ΔSV/BV remained unchanged from day 5 to day 15 in ZOL rats, while the NOM and OVX rats showed microdamage progression in ulnae (Fig. 4). In comparison, the OVX rats had more microdamage than the NOM rats despite lacking significant differences, which might be due to the higher standard deviation of ΔSV/BV on day 15 post-loading for the OVX. In fact, the maintained damage level in ZOL rats might be associated with efficient remodeling-based damage repair (Burr, 2014; Seref-Ferlengez et al., 2014; Seref-Ferlengez et al., 2015), while the increased damage level in NOM and OVX rats might be related to an excess amount of damage propagated relative to repair in the same time period (Wang & Niebur, 2006; Wu et al., 2013). Another possible explanation is that the increase of TMD, Ct.BV and Ct.Ar in the ZOL rats could have led to a reduced strain (rate) in the cortex while receiving the external mechanical loading, so that the critical stress intensity for crack initiation or propagation was not reached (Wright & Hayes, 1976; Zioupos, 2001). Therefore, the initial amount of microdamage might be determinant for their state of repair and propagation.

Past studies with similar experimental designs on rat forelimb showed that the microdamage volume ratio (SV/BV) was the highest 5 days post-loading (Zhang et al., 2018), and the damage density (Cr.Dn) reduced after 2 weeks of recovery post-loading (Herman et al., 2010). These findings are not exactly in line with the results in the present study, where the NOM and OVX rats had increased SV/BV 15 days post-loading. These discrepancies might be caused by several differences in experimental design such as rats’ gender, surgical treatments, and the methods used for microdamage quantification. A previous study using an ovariectomized ovine model revealed that the zoledronate treatment increased the average crack density but reduced average crack length based on histology analysis (Brennan et al., 2011). A study using an osteolytic rat model showed reduced SV/BV level due to zoledronate treatment, evaluated by contrast-enhanced micro-CT (Tolgyesi et al., 2022). As such, the reduced SV/BV level observed in the ZOL rats of the present study might have been caused by the reduced average length but not density, since the contrast enhanced micro-CT-based method is less sensitive than the histology-based method for damage detection. In general, the BPs treatment-related microdamage accumulation is more prominent in trabecular than in cortical bone (Allen & Burr, 2011).

Taken together, we conclude that the zoledronate treatment can reduce bone microdamage accumulation in cortical bone of osteopenic rats subjected to the same level of compressive loading. The lower accumulation of cortical microdamage might be caused by the improved structural properties in zoledronate treated rats, which might also determine the initial amount of loading-induced microdamage.

### 4.3 Microdamage accumulation correlates with periosteal bone formation

The correlation results in Table 4 show that more microdamage accumulation is associated with increased bone formation and decreased bone resorption on the periosteum of ulna. Based on visual observation in Fig. 3, the bone formed on the ulnar periosteum was mostly woven bone, which can help to restore the mechanical properties after tissue damage (Uthgenannt et al., 2007; Uthgenannt & Silva, 2007; Wohl et al., 2009). Moreover, the results show that the correlation between the microdamage level and the woven bone formation is stronger for the OVX and NOM rats than the ZOL rats, which might be caused by the lower microdamage levels in ZOL rats.

A past study with similar experimental design showed that zoledronate treatment led to higher periosteal bone formation compared to vehicle treatment (Feher et al., 2010). Although ZOL rats in our study showed periosteal bone formation, this response was weaker and less correlated with microdamage accumulation compared to the NOM and OVX rats. The contrasting results might be caused by several experimental differences, where in the Feher study, the *in vivo* mechanical loading was on a milder level (15 N, 360 cycles per day) for a longer period (1 week); the zoledronate was given as a single dose of 100 μg/kg; the ovariectomized animals were six months old, which might reveal some ageing-related characteristics of bone loss; and the dynamic histomorphometry for the evaluation of bone remodeling was based on histology. Nevertheless, we can confirm the cause and effect relationship between bone microdamage accumulation and periosteal bone formation (Uthgenannt et al., 2007), the extent of which was different between the normal, ovariectomized, and zoledronate-treated animals.

### 4.4 Limitations

In this study, an ovariectomized rat model was used to represent estrogen-deficiency related bone loss in human. One of the disadvantages of rat models for microdamage studies is that rats do not have baseline intracortical remodeling due to the lack of Haversian systems (Iwaniec & Turner, 2013; Jee & Yao, 2001; Lelovas et al., 2008). This is different in the human body, which makes it harder to translate rodent-based research to clinical settings (Allen, 2017). Rabbits are advantageous in this regard because they are considered to be the smallest laboratory animals with the capability of intracortical remodeling (Buettmann & Silva, 2016; Coates & Silva, 2020; Lelovas et al., 2008; Turner, 2001). Nonetheless, it is possible to initiate intracortical remodeling in rats with external stimulations such as mechanical loading (Bentolila et al., 1998). The age of the rats is also an important factor for bone loss. The ovariectomy-induced bone loss is often considered osteopenic rather than osteoporotic at 3 months old (Iwaniec & Turner, 2013; Jee & Yao, 2001; Kimmel et al., 2020; Lelovas et al., 2008), which is considered as a “mature rat model”, in contrast to the “aged rat model” of 6 to 12 months old (Kalu, 1991). Besides the advantages of shorter experimental duration and greater availability from animal suppliers, the 3 month old rat model is also meaningful to investigate the microdamage accumulation in young adult populations who undergo rigorous physical training (Bigosinski et al., 2010; Cole et al., 2020; Hod et al., 2006).

The analgesics used for the *in vivo* mechanical loading posed potential effects on bone metabolism. Although buprenorphine is usually considered a safe analgesic for laboratory animals (Guarnieri et al., 2012), clinical findings suggest that it can reduce bone density by altering the osteoblastic activity (Coluzzi et al., 2015). Meloxicam (metacam), a type of nonsteroidal anti-inflammatory drug (NSAIDs), can potentially inhibit osteoblastic activities thus reduce bone formation (Burch et al., 2021; Chang et al., 2005; Cottrell & O’Connor, 2010; Pountos et al., 2012). Though we still observed significant periosteal bone formation post-loading, these responses might have been stronger without the use of meloxicam. Nonetheless, all the rats received the same analgesic dosage in the present study to ensure meaningful between-group comparisons. The adverse effects of the analgesics were assumed to be negligeable.

The present study leveraged the non-destructive nature of the *in vivo* micro-CT imaging to evaluate bone (re)modeling. However, rats’ remodeling cycle is generally longer than the weekly scan interval (Baron et al., 1984; Jilka, 2013; Van et al., 1982; Weinstein et al., 1998). Therefore, the (re)modeling parameters obtained were not representative of a complete formation or resorption cycle, but rather local averaged rates including both modeling and remodeling events. Furthermore, the repeated anesthesia and immobilization associated with the *in vivo* imaging might have brought adverse effects to the animals’ well-being (Hohlbaum et al., 2017; Hotchkiss et al., 1998; Kádár et al., 2007; Kim et al., 2006; Patterson-Buckendahl et al., 2001; Stevens et al., 1975; Stollings et al., 2016). There is no direct evidence that the repeated irradiation performed caused significant bone loss to the scanned sites. Newer generations of micro-CT scanners might improve the image quality while reducing the scan duration, thus reducing the radiation doses. Deep learning algorithms also showed great potential for improving the interpretation of micro-CT scans (Wang et al., 2020).

In regard to the contrast-enhanced micro-CT imaging for microdamage evaluation, we used the same image resolution as for the *in vivo* scans but with higher voltage and current, longer exposure time, and more projections. This level of resolution was shown to be sufficient for detecting BaSO_4_ stained microdamage in rodent long bones (Turnbull et al., 2011) and human samples (Landrigan et al., 2011). Despite not being able to detect individual microdamages at this level of detail, the micro-CT-based SV/BV was proved to be well correlated with the histology-based measurement of microcrack density (Cr.Dn) (Landrigan et al., 2011; Zhang et al., 2018). Future studies can leverage the power of deep learning algorithms for microdamage detection in micro-CT images (Buccino et al., 2023; Caron et al., 2023).

Lastly, the process of BaSO_4_ staining might cause some alteration of the mechanical results. To account for this possibility, we conducted a preliminary test with 6 pairs of contralateral forelimb samples, where the results showed that none of the mechanical properties were significantly different between the contralateral ulnar samples. Therefore, we assumed that the between-group differences were not significantly changed by staining since all the samples went through the same processing steps. We considered that the advantages of reusing the same animals for mechanical testing outweighed their disadvantages based on the 3R principles (Sneddon et al., 2017).

## 5 Conclusion

In the present study, we developed and exploited an *in vivo* experimental platform combining the evaluation of bone microdamage, (re)modeling, mechanical properties as well as their correlations in the same cohort of animals receiving different treatments. Based on our results and evidence from the literature, we conclude that zoledronate treatment can reduce cortical microdamage accumulation and improve osteopenic conditions of ovariectomized rats, which might be explained by the enhancement of several bone structural properties. Microdamage accumulation showed positive correlations with periosteal bone formation, while exhibiting negative correlations with bone resorption and mechanical properties. The extent of these correlations is different in normal, ovariectomized and zoledronate-treated animals. Our results provide insights towards further investigation of pharmaceutical treatments for osteoporosis and osteopenia. The same experimental concept can be applied in future studies on microdamage and drug testing.

## Acknowledgements

This study was made possible by a micro-CT platform supported by the TransMedTech Institute. The authors would like to thank Dr. Ryan K. Roeder, Dr. Cari M. Whyne, Dr. Da Jing, and their teams for sharing the BaSO_4_ staining procedure. Special thanks to Dr. Bettina Willie and her team for sharing the bone (re)modeling evaluation algorithm.

