## Supplementary Material for "Zoledronate reduces loading-induced microdamage in cortical ulna of ovariectomized rats"

### Analysis of trabecular tibia

The right hindlimb (proximal metaphysis) of all the rats were scanned by micro-CT on week 2 and week 12 to confirm the effects of ovariectomy and zoledronate treatment. The scan settings were the same as explained in section 2.2 of the article. The proximal tibial VOIs of 2 mm long were taken starting from 2 mm distal to the proximal growth plate. The trabecular VOIs were then segmented using a semi-automatic selection tool in CT Analyzer. A global threshold of 80/255 (0.49 g/cm$^3$) was set for the following trabecular analysis, which was determined by an automatic thresholding function in CT Analyzer and visually verified in the images and histograms. The following trabecular parameters were analyzed: bone mineral density (BMD, g/cm$^3$), trabecular bone volume (Tb.BV, mm$^3$), trabecular bone surface area (Tb.BS, mm$^2$), trabecular bone volume fraction (Tb.BV/TV), trabecular thickness (Tb.Th, μm), trabecular number (Tb.N, 1/mm), trabecular separation (Tb.Sp, μm), and connectivity density (Conn.Dn, 1/mm$^3$).

Supplementary Table. Tibial dynamic histomorphometry (n=16/group). "↑" or "↓" refers to a significant increase or decrease compared to week 2. "N", "O", or "A" refers to a significant difference compared to the NOM group, the OVX group, or all the other groups. Bonferroni correction was applied for the *p*-value calculation in the *post hoc* tests and multiple comparisons, after which the statistical significance level (alpha) was set to 0.05. The cells of statistical significance are in bold.

| Time | Group | BMD (g/cm$^3$) | | Tb.BV (mm$^3$) | | Tb.BS (mm$^2$) | | Tb.BV/TV (1) | |
|---|---|---|---|---|---|---|---|---|---|
| Week 2 | NOM | **0.44 ± 0.05** | **A** | 9.26 ± 1.13 | | 555.12 ± 75.28 | | **45.15% ± 8.83%** | **A** |
| | OVX | **0.28 ± 0.04** | **A** | **5.16 ± 1.44** | **N** | 460.73 ± 122.69 | | **21.95% ± 5.59%** | **A** |
| | ZOL | **0.38 ± 0.03** | **A** | **8.24 ± 1.23** | **O** | **630.64 ± 124.22** | **O** | **35.11% ± 4.88%** | **A** |
| Week 12 | NOM | **0.47 ± 0.05** | **A** | **10.15 ± 1.21** | **A** | **548.31 ± 76.30** | **A** | **49.83% ± 6.65%** | **A** |
| | OVX | **0.22 ± 0.03** | **A↓** | **3.26 ± 0.68** | **A↓** | **233.60 ± 57.50** | **A↓** | **17.71% ± 2.48%** | **A** |
| | ZOL | **0.66 ± 0.06** | **A↑** | **22.57 ± 5.77** | **A↑** | **704.38 ± 109.81** | **A** | **73.09% ± 7.74%** | **A↑** |

| Time | Group | Tb.Th (μm) | | Tb.N (1/mm) | | Tb.Sp (μm) | | Conn.Dn (1/mm$^3$) | |
|---|---|---|---|---|---|---|---|---|---|
| Week 2 | NOM | **67.73 ± 7.42** | **A** | 6.63 ± 0.79 | | 88.52 ± 9.68 | | 2579.78 ± 350.05 | |
| | OVX | **49.37 ± 6.04** | **A** | **4.46 ± 1.08** | **N** | **131.10 ± 30.91** | **N** | 1700.91 ± 579.30 | |
| | ZOL | **55.82 ± 5.56** | **A** | **6.29 ± 0.88** | **O** | **94.82 ± 11.31** | **O** | **2661.36 ± 648.74** | **O** |
| Week 12 | NOM | 73.27 ± 7.14 | | **6.80 ± 0.58** | **A** | **84.80 ± 6.56** | **A** | 2607.99 ± 421.45 | |
| | OVX | **72.14 ± 6.68** | ↑ | **2.48 ± 0.49** | **A↓** | **153.97 ± 14.28** | **A** | **949.94 ± 297.08** | **A↓** |
| | ZOL | **89.96 ± 10.59** | **NO↑** | **8.17 ± 0.75** | **A↑** | **67.95 ± 8.46** | **A↓** | 2482.04 ± 738.30 | |
